# Design of a novel DNA Gyrase B inhibitor with a rhodanine scaffold: *in silico* and *in vitro* approaches

**DOI:** 10.1101/2021.01.24.428017

**Authors:** Akhila Pudipeddi, Sahana Vasudevan, Karthi Shanmugam, Alex Stanley, Vairaprakash Pothiappan, Prasanna Neelakantan, Adline Princy Solomon

## Abstract

Methicillin-resistant *Staphylococcus aureus* (MRSA) and vancomycin intermediate-resistant *Staphylococcus aureus* (VRSA) is one among the WHO high priority pathogens. Among these two, MRSA is the most globally documented pathogen that necessitates the pressing demand for new classes of anti-MRSA drugs. Bacterial gyrase targeted therapeutics are unique strategies to overcome cross-resistance as they are present only in bacteria and absent in higher eukaryotes. The GyrB subunit is essential for the catalytic functions of the bacterial enzyme DNA Gyrase, thereby constituting a promising druggable target. The current study performed a structure-based virtual screening to designing GyrB target-specific candidate molecules. The *de novo* ligand design of novel hit molecules was performed using a rhodanine scaffold. Through a systematic *in silico* screening process, the hit molecules were screened for their synthetic accessibility, drug likeliness and pharmacokinetics properties in addition to its target specific interactions. Of the total 374 hit molecules obtained through *de novo* ligand design, qsl-304 emerged as the most promising ligand. qsl-304 was synthesized through a one-step chemical synthesis procedure, and the *in vitro* activity was proven, with an IC_50_ of 31.23 μg/mL against the novobiocin resistant clinical isolate of *Staphylococcus aureus sa*-P2003. Further studies on time-kill kinetics showed the bacteriostatic nature with the diminished recurrence of resistance.

## Introduction

Multidrug resistance is a major global public health issue caused due to the indiscriminate use of broad-spectrum antimicrobial and also the usage of counterfeit antimicrobials *(1)*. Such falsified and substandard antimicrobials may contain toxic doses of dangerous ingredients, compromise the treatment of chronic and infectious diseases, causing disease progression, drug resistance, and increases the mortality rate *(2).* Recent hospital-based surveillance reports suggest the emergence of a resistant group of nosocomial pathogens comprising *Enterococcus faecium, Staphylococcus aureus, Klebsiella pneumoniae, Acinetobacter baumannii, Pseudomonas aeruginosa,* and *Enterobacter* species (ESKAPE pathogens) that have diminished the efficacy of many frontline antibacterial agents, which will result in a global financial burden of ~100 trillion USD by 2050 *(3–6)*.

According to the Centre for Disease Control and Prevention (CDC), one of the ESKAPE pathogens, the Methicillin resistant-*S. aureus* is a crucial player to cause severe infection both in community-acquired and hospital-related infections *(7,8).* The widespread usage of antibiotics, acquisition and accumulation of resistance-conferring alleles have resulted in the emergence of MRSA strains to develop cross-resistance to vancomycin (VAN), trimethoprim-sulfamethoxazole, β-lactams, tetracycline, clindamycin, quinolones, and aminoglycosides *(9,10,6).* The resistance of MRSA to already existing antibacterial drugs might be due to the Staphylococcal Chromosomal Cassette gene mec (SCCmec) and Panton-Valentine Leukocidine (PVL) toxin gene that enhances bacterial virulence *(11)*.

To avoid cross-resistance with currently used antibacterial drug classes and meet the inadequacies of current anti-MRSA therapies, there is a pressing demand for new classes of anti-MRSA drugs. Indeed, the need for designing anti-MRSA drugs may discover novel mechanisms of action or focus on previously unexploited binding sites on existing targets or develop novel scaffolds without developing cross-resistance *(12,13).* In this accord, bacterial gyrase targeted therapeutics is unique since it is the universal enzyme required for bacterial survival and is absent in higher eukaryotes*(14)*. DNA gyrase (gyrase), a member of bacterial topoisomerases is known to control the DNA dependent processes by introducing transient breaks to both DNA strands *(15)*, in addition to relieving the torsional tension, by introducing negative supercoils to the DNA molecule *(16).* DNA gyrase is a heterotetrameric protein composed of two GyrA subunits where the DNA cleavage site is located, and two GyrB subunits that provide the energy necessary for the catalytic function of the enzyme through ATP hydrolysis *(4)*. Therefore, drugs that target bacterial topoisomerases act by two main mechanisms, either by stabilizing the complex between the DNA molecule and the GyrA active site of the enzyme (e.g. quinolones) or by inhibiting the ATPase activity of the GyrB subunit (e.g. aminocoumarin class of inhibitors) *(17).*

Novobiocin, a representative of the natural aminocoumarins, was the only clinically-approved potent inhibitor of GyrB with inhibition (K_i_) and binding (K_d_) constants in the low nanomolar range (7–15 nM) *(18),* until recently, wherein the United States Food and Drug Administration (FDA) withdrew its approval due to issues of toxicity and poor physiochemical properties, urging the discovery of an improved successor to novobiocin *(19).* The discovery of new drugs with novel scaffolds targeting the ATP binding site of GyrB without any cross-resistance and toxicity issues will complement the failure of novobiocin *(17).* Several reports suggest that heterocyclic compounds with rhodanine scaffold are known to inhibit several pharmaceutical targets from demonstrating enhanced antimicrobial activity *(20).* 3, 5-disubstituted rhodanine derivatives inhibited the S. aureus supercoiling activity by targeting the GyrB ATPase function*(21)* and other rhodanine derivatives such as 5-(2-hydroxybenzylidene) has shown moderate antibacterial activity, reported as GyrB inhibitor *(22)*, phenylalaninederived (z)-5-arylmethylidene rhodamine *(23)*,2-(5-(3,4-Dichlorobenzylidene)-4-oxo-2-thioxothiazolidin-3-yl)-3-phenylpropanoic acid and 2-(5-(3-Phenoxybenzylidene)-4-oxo-2-thioxothiazolidin-3-yl)-3-phenylpropanoic acid *(24)* have shown significant anti MRSA activity, can be used for treatment of gram-positive MRSA *(25).* Rhodanines which have a privileged scaffold in drug discovery, wide spectrum of pharmacological activity *(26)* and structural modification enable potent, selective drugs to be developed.

The current study is aimed to identify potent compounds that target the GyrB ATP binding site through a systematic virtual screening process. In this regard, we have extended our search to find efficient GyrB-targeted inhibitors using *de novo*-based ligand design. As a lead to progress, an insight of recent work on NMR-fragment-based binding study against the co-crystallized full-length *S. aureus* GyrB ligand *(27),* was taken as a reference to optimize the target-specific hit molecules with rhodanine scaffold using LigBuilder. The bioavailability and pharmacokinetics of the top-hit ligands were further scored based on ADMET properties, and we then evaluated the biological efficacy against MRSA *in vitro*.

## Materials and methods

### Protein Selection

The current computational studies were done with the crystal structure of *S. aureus* GyrB complexed with a fragment (indolinone derivative), PDB ID: 5CPH. There is a total of 32 structures deposited for *S. aureus* GyrB in RCBS Database (www.rcbs.org). Among them, the structure with PDB ID – 5CPH has the best resolution of 1.2 Å. The considered protein, GyrB consists of two chains (homodimer), A and B having 212 amino acids. For further studies, chain A of GyrB was chosen.

### *De novo* Ligand Design

A structure-based drug design methodology termed *de novo* ligand design was carried out using LigBuilder *(28). De novo* design is a successful ligand design method where the novel ligands are built in a stepwise manner considering the protein binding pocket constraints. The unique advantage of this method is the generation of molecules based on the target protein, which is unique and not available in any other small molecule databases. LigBuilder is one of the acclaimed *de novo* tools which builds the ligands from the library of organic molecules having 60 fragments valuable to the drug design process. There are two modules of “growing” and “linking” to build ligands which are controlled by genetic algorithms. Among the two approaches, the current study was designed based on the “grow” concept. Here, the seed structure plays an important role and is the starting point of the whole process of *de novo* design of hit optimization. Rhodanine scaffold was designated as the seed structure for the hit optimization process. All the molecules generated out of the *de novo* drug design will have the structural framework of rhodanine. Another notable requirement is the pre-docked protein-ligand complex for the program to locate the binding pocket. In our case, as there was no co-crystalized rhodamine-GyrB complex structure in the protein data bank, we docked rhodanine in the binding pocket of S. *aureus* GyrB (PDB ID: 5CPH) and used it for pocket exploration. The “building” of the molecule from the rhodanine core was from the two hydrogen atoms at the C5 position. The next subsequent step is the mutation operation where the generated molecules are improved for their ligand-binding affinity. The building up process is further amplified using genetic algorithms. Finally, *de novo* designed molecules are obtained.

### Synthetic Accessibility Score

One of the significant limitations of the *de novo* ligand design is the synthetic feasibility. Thus, to overcome this barrier, the newly developed *de novo* compounds were subject to its synthetic accessibility. SYBA (SYnthetic Bayesian Accessibility) was used to classify the *de novo* designed molecules as easy – (ES) and hard – to – synthesize (HS) organic compounds based on the analysis of the individual substructures of the given hit molecules *(29).* SYBA is the latest improved version of SAScore and SCScore with ES molecules trained with the molecules of ZINC15 database and HS molecules from Nonpher methodology. SYBA works on Bernoulli naïve Bayes classifier where each compound is fragmented into substructures, and each fragment is scored. The individual substructures score is added to get final SYBA score. The analysis is based on the score obtained by Bayesian analysis. The positive scores imply ES and negative scores, HS. SYBA is publicly available at https://github.com/lich-uct/syba under the GNU General Public License.

### Protein preparation

Schrödinger’s Protein Preparation Wizard *(30)* is used for the preparation of three-dimensional structure of ATP binding domain of *S. aureus* GyrB complexed with a (3E)-3-(pyridin-3-ylmethylidene)-1,3-dihydro-2H-indol-2-one inhibitor (PDB ID: 5CPH). The protein preparation workflow briefly includes the addition of missing hydrogen atoms, enumerate bond orders to HET groups, removing co-crystallized water molecule (non-structural), determining most likely protonation state, correcting potentially transposed heavy atoms, and optimizing proteins hydrogen bond network. Finally, the protein was subjected for a restrained minimization that allows hydrogen atoms to be freely minimized, while allowing for sufficient heavy atom movement to relax strained bonds, angles and clashes.

### Receptor grid generation

The recognition of the active site or the binding site of the target structure accurately is a key step in drug designing through computational docking of the novel chemical moieties. Schrödinger GLIDE (Grid based ligand docking with energetics) docking protocol generates a grid around the active site of the protein for ligand docking *(31)*. The shape and properties of the receptor are represented on a grid by several different sets of fields that provide progressively more accurate scoring of the ligand poses. The grid box was generated using receptor grid generation panel. Grid box is positioned on the centroid of the co-crystallized ATP binding domain inhibitor (3E)-3-(pyridin-3-ylmethylidene)-1,3-dihydro-2H-indol-2-one. As the ATP binding pocket of GyrB is well characterized and reported, we manually checked and ensured that the generated grid encloses all the important amino acids of GyrB ATP binding site (D81, R84, G50, R144 and HOH427). The atoms of protein were fixed within the default parameters of the radii of Vander Waal’s scaling factor of 1 Å with partial charge cut-off of 0.25 Å using OPLS3e force field and 20 Å docked ligand length. The dimensions of the grid box and receptor setup were x = 22 Å, y = 22 Å, z = 22 Å and x = 64 Å, y = 64 Å, z = 64 Å during docking study, respectively, with a grid space of 1 Å.

### Ligands preparation

An accurate 2D to 3D conversion of ligand molecules is a key precursor to computational docking studies. The 374 novel ligands designed through LigBuilder algorithm was prepared using LigPrep module implemented in Schrödinger (Schrödinger Release 2020-4: LigPrep, Schrödinger, LLC, New York, NY, 2020.). LigPrep generates accurate, energy minimized 3D molecular structure using OPLS3e force field. It also applies sophisticated rules to correct Lewis structures and eliminate mistakes in ligands to reduce downstream computational errors. LigPrep expands tautomeric and ionization states, ring conformations, and stereoisomers to produce structural diversity from a single input structure.

### High Throughput Virtual Screening and Docking

The prepared 374 molecules generated by LigBuilder was taken further for high throughput virtual screening (HTVS) against a target protein (PDB ID: 5CPH). The ligands were docked using Glide (Flexible mode) to evaluate the binding modes. Glide extra precision docking was carried out for all the 374 molecules *(31).*

### Calculation of binding free energies by prime MM-GBSA approach

The ligands and receptor complexes were analyzed for their binding free energies by prime MM-GBSA (Molecular Mechanics Generalized Born Surface Area) module of Schrödinger suite with the OPLS3e force field *(32).* The prime MM-GBSA approach is used to calculate ΔG bind of each ligand which is based on the docking complex interactions. The binding free energy was calculated using the following equations.

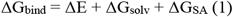

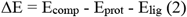

Where Ecomp, Eprot and Elig are the minimized energies found for the protein-ligand complex, protein and ligand molecule respectively.

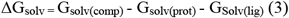

Where ΔG_solv_ is the born electrostatic solvation energy found for the complex and ΔG_SA_ is the non-polar contribution to solvation energy by the surface area.

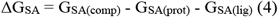

Where GSA_(comp)_, GSA_(prot)_, and GSA_(lig)_ are the surface area energies for the protein ligand complex, protein and ligand respectively.

### Lipinski’s and Veber’s Rules

Schrödinger Qikprop (Schrödinger, LLC, New York, NY, 2020) was used for a further level of filtering to identify the molecules having drug-likeliness using Lipinski’s Rule of Five (Hydrophobicity less than 5, molecular weight less than 500 Da, hydrogen bond (HB) donor and acceptor less than 5 and 10 respectively). In addition to that the hit molecules were assessed for Veber’s rule having rotatable bonds <10 and polar surface area < 140 Å were also taken for the screening process.

### ADMET Prediction

The molecules which did not have any violation of both Lipinski’s and Veber’s rule were further assessed for the ADMET properties using Discovery Studio 2.5. ADME (Absorption, Distribution, Metabolism and Elimination) assessment is required to filter the hit molecules with desirable pharmacokinetic properties. The ADME descriptors such as aqueous solubility, Blood-brain barrier penetration, Cytochrome P450 2d6 inhibition, Hepatotoxicity, Human Intestinal absorption, and Plasma protein binding. The hit molecule(s) which satisfies the above properties are taken for *in vitro* analyses.

### Synthesis of (Z)-(5-((4-oxo-2-thioxothiazolidin-5-ylidene)methyl)furan-2-yl)methyl acetate

A suspension of rhodanine (18 mmol, 2.34 g) and 5-hydroxymethylfurfural (19.4 mmol, 2.45 g) in glacial acetic acid (19.4 mL) was heated to 80 °C for complete dissolution *(33).* The resulting solution was treated with sodium acetate (77.8 mmol, 6.37 g) at 80 °C for 15 h. Completion of the reaction was monitored by TLC analysis using a solvent mixture containing hexane and ethyl acetate (1:1, v/v) as an eluent. Then the reaction mixture was treated with water (50 mL) and extracted using ethyl acetate (3 x 30 mL). The combined organic extract was dried over sodium sulphate and filtered. The solvent was removed by rotary evaporation and the resulting residue was purified by column chromatography [silica, ethyl acetate: hexane (1:9)] to obtain the compound (Z)-(5-((4-oxo-2-thioxothiazolidin-5-ylidene)methyl)furan-2-yl)methylacetate (indicated as qsl-304, herein) as a yellow solid. 1H-NMR spectroscopic analysis was used to confirm the synthesized compound.

### Antibiotic and Test Compound Preparation

The antibiotic novobiocin was obtained as a DMSO-soluble powder with 96% purity (Sigma-Aldrich (St Louis, MO, USA). The stock solution of the test compound, qsl-304 was prepared in 1% DMSO. The final concentration of DMSO was maintained as <0.5%.

### Bacterial Strains and Culture Maintenance

The reference strain, *S. aureus* ATCC43300 was purchased from the American Type Culture Collection (ATCC; Manassas, VA) and the clinical isolates, *Sa*-P1934, *Sa*-P1920, *Sa*-P1996, *Sa*-P2003, *Sa*-2052, *Sa*-AB77, *Sa*-AB459, *Sa*-AB472 were received from JSS Medical College, Mysore, India. The cultures were maintained in as glycerol stock at −80°C.

### Novobiocin Susceptibility Testing

Susceptibility to novobiocin was performed for both the standard and clinical isolates by the broth microdilution method according to the CLSI guidelines *(34).* The antibiotic novobiocin was obtained as DMSO-soluble powder (Sigma-Aldrich (St Louis, MO, USA) and used for MIC determinations as interpreted in the interpretive susceptibility criteria reported in the appropriate CLSI Tables (M100-S25, CLSI, 2015).

### Antibacterial evaluation of qsl-304

The doubling dilutions of the test compound qsl-304 were serially aliquoted into each well of a 96-well microplate using 8-points, covering a range from 500-3.9 μg/mL as to reach final assay volume of 100μL/well. Then, 10μL culture suspension of the exponentially growing cells with a seed density of 10^5^ CFU/ml was added into each well. The solvent control (medium with DMSO) and negative controls (medium with culture inoculum; medium with broth) were maintained in all experiments. All independent culture conditions were performed in triplicate. Further, the plates were incubated at 37°C for 24 h in ambient air and growth was assessed at OD595 using microplate reader (iMark, Bio-Rad, Japan). The lowest concentration that completely prevented bacterial growth was categorized as MIC.

### Time-dependent Kill Assay

The time-dependent effect of qsl-304 on the growth of the novobiocin resistant clinical isolate *sa-* P2003 was performed by the broth macro-dilution method as per the CLSI guidelines. The exponentially growing cells were used as bacterial inoculum with a seed size of 10^5^ CFU/ml. The test compound qsl-304 was added to 10ml of the inoculum suspension to achieve a final concentration of 0.5xMIC, 1xMIC, 2xMIC, 4xMIC, 8xMIC and 16xMIC and maintained at 37°C in ambient air for 24 h. A growth control (bacterial inoculum without the test compound) was maintained in all independent trials. After 24h, the aliquots from the inoculum culture were removed at defined time-points (0, 2,4,6,8,10 and 24 h) and ten-fold serial diluted in saline and the growth was determined by measuring the absorbance at OD600 and compared with that of the control (growth media with *Sa*-P2003).

The data was analyzed by evaluating the number of viable cells that yielded *Δ*(log10 CFU/mL) values of −1 (90% killing), −2 (99% killing), and −3 (99.9% killing) at various time-point intervals compared to viable counts at 0 h. Meanwhile, the bactericidal activity was calculated as a reduction of at least 99.9% (≥3 log10) of the total count of CFU/mL in the original inoculum *(35).*

### Frequency of Spontaneous Resistance to qsl-304

The bacterial strains *Sa*-P2003 and ATCC43300 were cultured in CA-MHB and MHA to examine the frequency of spontaneous resistance to a higher concentration of qsl-304. Briefly, bacterial cultures that reached an exponential phase in CA-MHB were serial diluted to a seed inoculum size of ~10^9^ cfu/mL and plated onto agar containing 0.5x, 1x, 2x and 4x MIC of the test compound to calculate the generation and enumeration of resistant mutants. The cultures were also diluted in PBS and dilutions containing ~10^2^ cfu/mL were plated onto drug-free agar for the enumeration of total viable cells. Plates were incubated at 37°C for 48 h. The experiment was carried out in triplicates and the negative control (media without qsl-304) was maintained *(36,37).* After the stipulated time (48h), the viable cells were counted and the frequency of spontaneous resistance towards the action of qsl-304 was calculated using the following formulae:

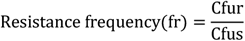

Cfur = CFU/mL of test strain in the presence of qsl-304

Cfus= CFU/mL of test strain grown in the absence of qsl-304

### Data Analysis

All the *in vitro* experiments were conducted in biological and technical replicates. All the data obtained were analyzed using GraphPad Prism 8.0.2 software. Dose-response curves were plotted by fitting the inhibition data using a 4-parameter logistic equation.

## Results and Discussion

### Design and Synthesis

The advent of computer-aided drug design reduces the traditional drug discovery process. *De novo* ligand design is one such structure-based virtual screening (SBVS) approach on generating a set of target-specific novel chemical hits with desirable activity – enriched profiles. In this regard, the current study aimed to identify potential GyrB inhibitors through a systematic virtual screening process (Figure 1).

**Figure 1:**
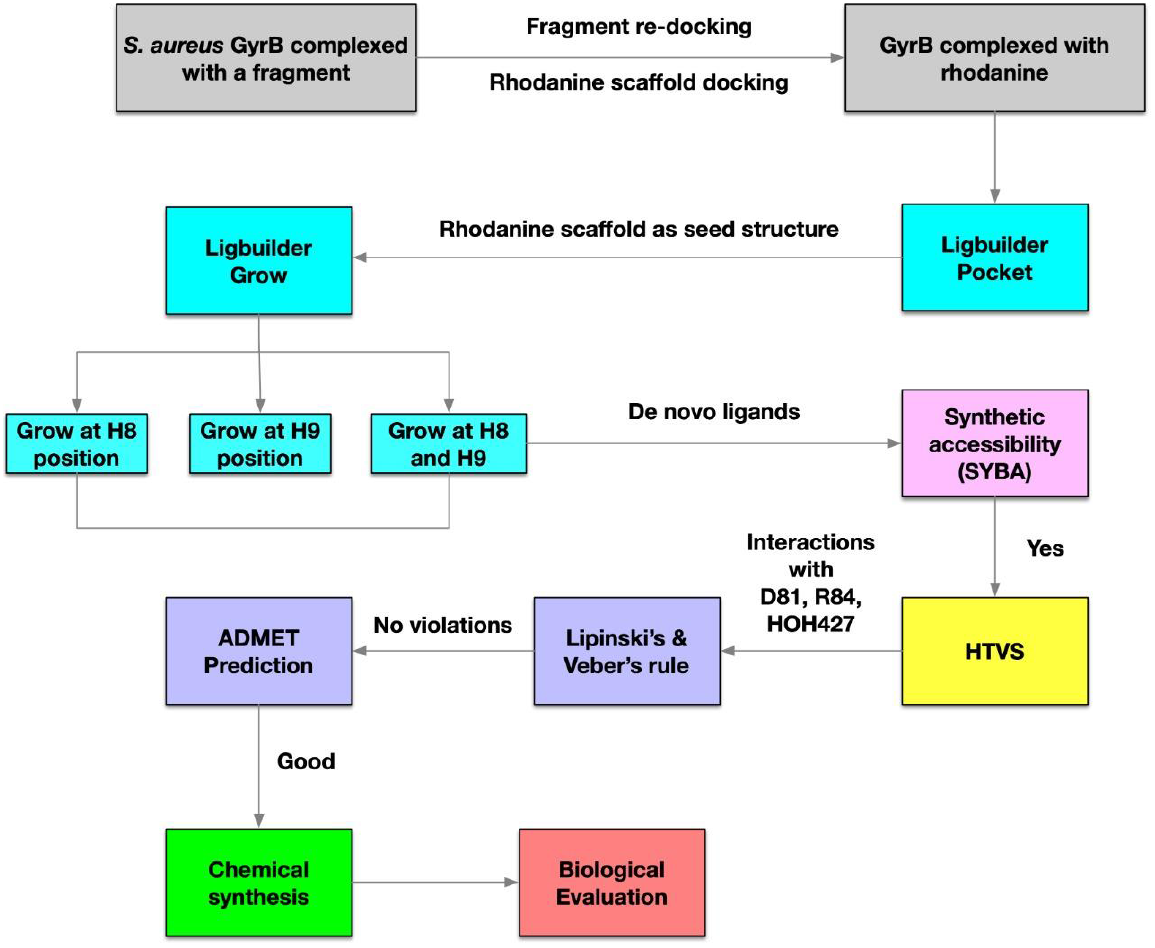
In silico-based hit optimization workflow

The target protein, protein-ligand interaction complex, and seed structure(s) are the cornerstones that aid in the successful hit optimization process. As mentioned earlier, the target protein is GyrB protein co-crystallized with indolinone fragment (PDB ID:5CPH). The indolinone fragment interacts with the GyrB in its ATP binding site (Asp81 and a crystallographically conserved water molecule) similar to the other potent inhibitors *(27)*. Thus, this crystal structure was taken for the further docking and ligand building processes. Furthermore, the docking algorithm was validated with a molecular redocking procedure. Indolinone was separated from the complex structure and redocked into GyrB. The glide score of the redocked complex was –5.68 with the RMSD of 1.23Å between the crystal and computed docked pose of the indolinone fragment. As shown in Figure 2a, binding interaction analysis of the re-docked complex revealed that the N-H group at position 1 and C=O at position 2 participated in the hydrogen bond network with the carboxylate side chain of Asp81 and crystallographic conserved water molecule respectively. These results were in coherence with the crystal pose and hence suggested the consistency of the docking algorithm and force fields used.

**Figure 2:**
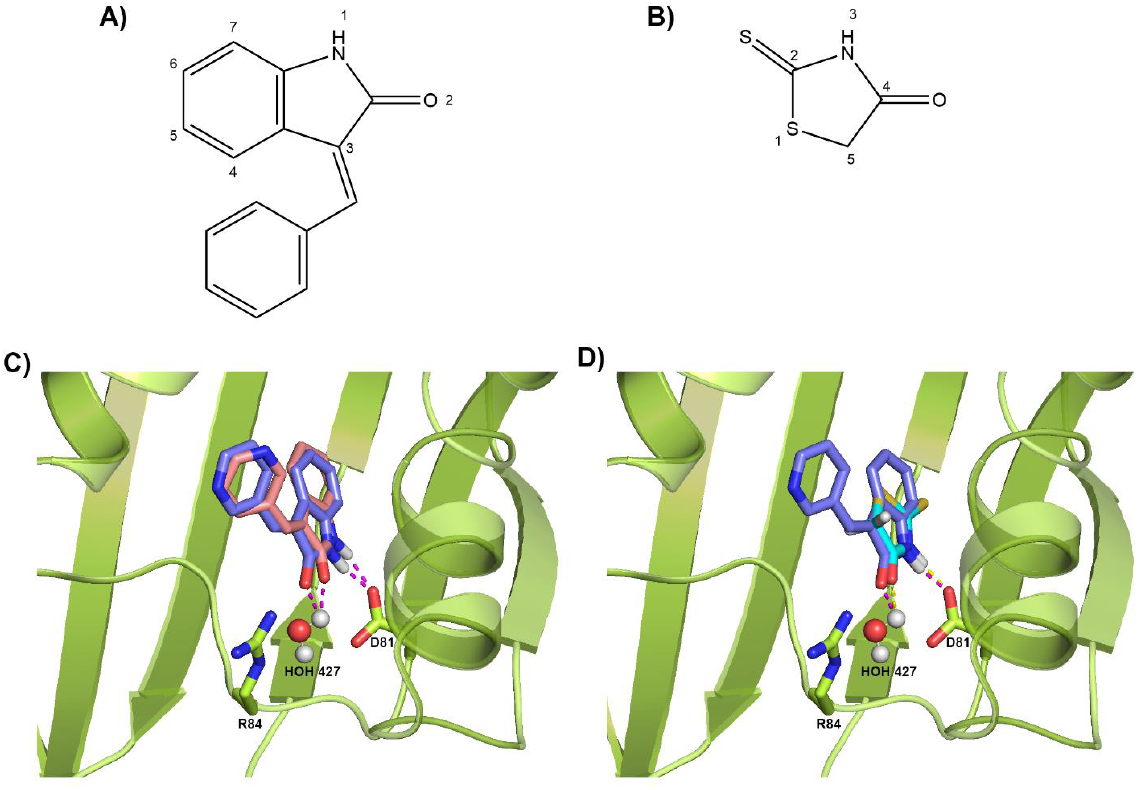
A) Structure of the Indolinone fragment B) Structure of rhodanine scaffold used in this study. C) Illustration of redocking of co-crystallized indolinone fragment with GyrB (PDB ID:5CPH). GyrB is represented as a green ribbon. Crystal and docked poses of indolinone were represented as blue and salmon sticks. Hydrogen bonding interactions with Asp 81 and the crystallographically conserved water (427) were designated as pink dotted lines. D) Docking of rhodanine fragment on to GyrB. The green ribbon represents GyrB protein structure. Blue stick represents crystal pose of indolinone. Cyan stick represents the docked pose of rhodanine. The pink and yellow dotted line represents the hydrogen bonding interactions made by indolinone and rhodanine fragment, respectively.

After the validation of the docking algorithm, seed structure selection was performed. Seed structure is an integral component of the *de novo* drug design process as it is the starting point of the generation of novel hit molecules. For the current study, rhodanine moiety (2 – thioxothiazolidin – 4 -one) was taken as the building block for the hit generation. Considered to be the “Privileged Scaffold”, rhodanine belongs to the family of five-membered multi-heterocycles (FMMH) having an exocyclic sulphur atom *(26)* and owing to its distinct intermolecular interaction pattern, rhodanine scaffold feature in the numerous pharmaceutical molecules *(38)*. Especially in the post-antibiotic age of antimicrobial resistance, several hit molecules having rhodanine scaffold were proved to have effective antimicrobial activity against susceptible as well as MDR strains of both Gram-negative and Gram-positive bacteria *(20)*. Therefore, the present study was directed towards the development of novel hit molecules from rhodanine against *S. aureus* GyrB. Before proceeding with the “growth” of the seed structure, *de novo* ligand design approach uses protein-ligand complex information as the input to construct and design novel hit molecules. Even though the crystal structures of hit molecules having rhodanine scaffold are available *(21,39)*, interestingly, there was no experimentally solved structure of GyrB directly complexed with rhodanine. Hence, to prepare the initial complex structure for *de novo* ligand design, we docked the rhodanine moiety into GyrB (PDB: 5CPH) and used it for further hit development and optimization. The docking result shows that the N-H at position three and C=O at position 4 of rhodanine were participating in hydrogen bonding interactions with the side chain of Asp81 and a crystallographic conserved water molecule, respectively (Figure 2b).

The GyrB-rhodanine docked complex was provided as input for LigBuilder. The binding pocket identification and the pharmacophore elements of the input complex structure were identified by running the pocket module of LigBuilder. LigBuider offers two different strategies (grow and link) to develop novel hit molecules, and in this study, we utilized the grow approach. Two positions of the rhodanine scaffold (N3 and C5) were extensively explored for the development of potent antimicrobial derivatives *(21,39,20).* In our case, the N3 position of rhodanine participates in the hydrogen bonding interaction with the Asp81, and hence the two hydrogen atoms at C5 position were considered for growing the fragment (Figure 3). To obtain structurally diverse rhodanine derivatives, the two hydrogens at C5 positions were grown separately and simultaneously. LigBuilder generated a total of 374 novel hit molecules with the rhodanine scaffold.

**Figure 3:**
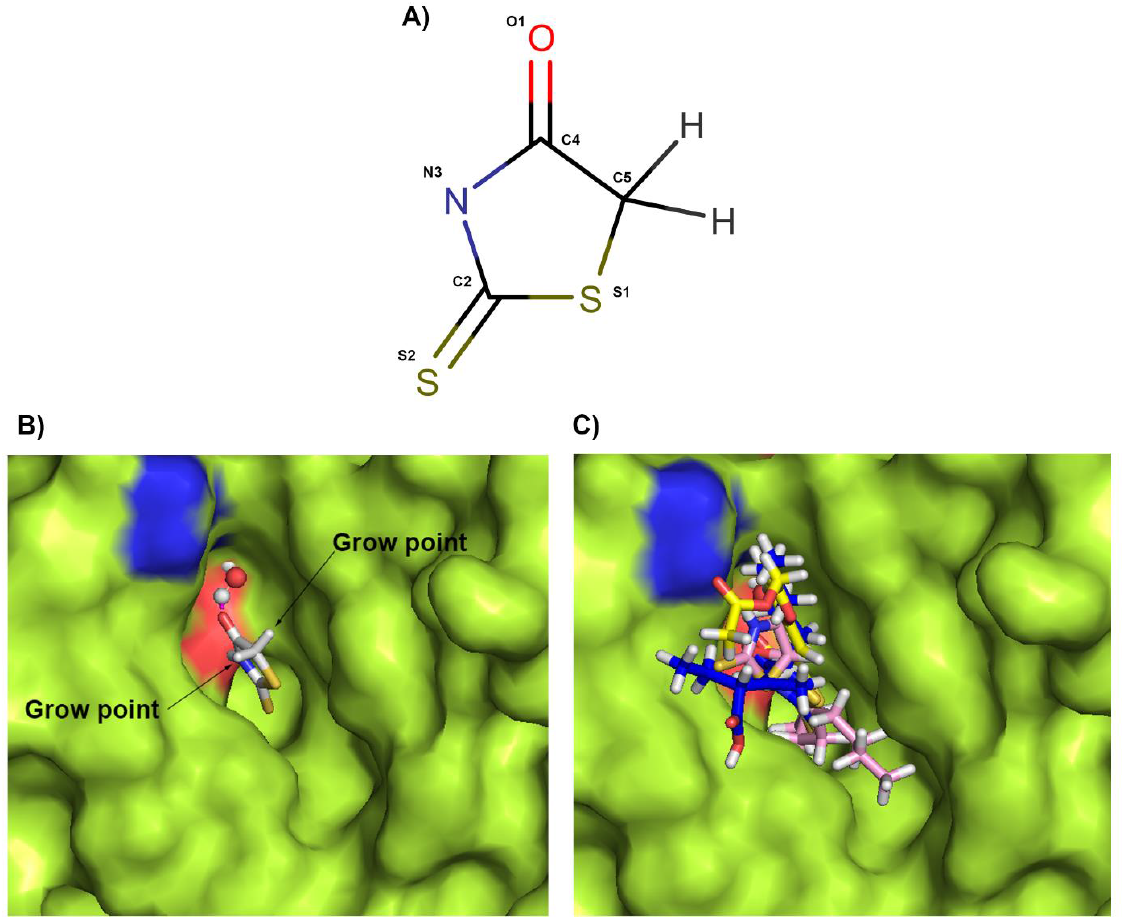
Sketch map of the growing operation. (A) Seed Structure (rhodanine scaffold) (B) The docked pose of rhodanine (seed structure) in the ATP binding domain of GyrB. The two hydrogen atoms at C5 positions were labelled as growing points. (B) Representation of molecules grown at two hydrogens at C5 positions of rhodanine individually and simultaneously. The newly developed ligands were represented as sticks.

Despite the fact that synthetic accessibility is integrated in the *de novo* ligand design program generating a chemical score, the molecular complexity results in insufficiencies *(40,41),* encouraging us to employ a fragment-based scoring method, SYBA, to evaluate the synthetic accessibility of the generated molecules *(29).* The SYBA score is assigned to the molecules derived from the ECFP8 fragments randomly present in the ZINC15 database for ES molecules and Nonpher approach generated compounds for HS molecules. Since the number of stereo centres is limited, the compounds having multiple stereo centres are penalized. Hence, the prediction confidence is expressed as a positive SYBA score denotes ES, and a negative score indicates HS. Based on this, the 374 molecules were analyzed and assigned an SYBA score. The hit molecules having positive SYBA score were filtered and labelled as ES. Through this filtration process, 374 molecules were reduced to 58 molecules. To further improve the stringency of the hit optimization process, these molecules were looked for the key molecular interactions (D81, R84 and HOH427). Among the two pharmacophore features of the previous established GyrB inhibitors, hydrogen bond – acceptor motif is given primary importance for the successful inhibition of the GyrB ATP activity. The second feature is the presence of an aromatic ring forming cation- π interaction with Arg-84 and a hydrogen bond with Arg −144 side chain *(27)*. This process reduced the number of molecules from 57 to 44 hit molecules.

Finally, the drug-likeness and the pharmacokinetics property were evaluated for the 44 molecules. Lipinski’s Rule of Five *(42)* and Veber’s rule *(43)* were assessed for the 44 hit molecules, to screen the “druggable” compounds specifically from the oral bioavailability viewpoint. Table 1 lists the Lipinski’ s and Veber’s parameters of the eight molecules which followed both Lipinski’s and Veber’s rule. Out of the 44 hit molecules which can be easily synthesized, only 8 obeyed Lipinski’s rule of five and Veber rule. The 2D interaction map of the eight molecules are given as supplementary figure (Figure S1). For the eight molecules, logP was within the preferred range of <5, and the molecular weight was ≤500 Da depicting good oral bioavailability as well as membrane permeability. Also, Veber’s rule of a number of rotatable bonds were ≤ 10 for all the eight hit molecules and PSA ≤ 140 Å showcasing good intestinal availability. Stereoselectivity of the hit molecules for optimal target binding was evaluated with the number of rotatable bonds *(43)*

**Table 1:**
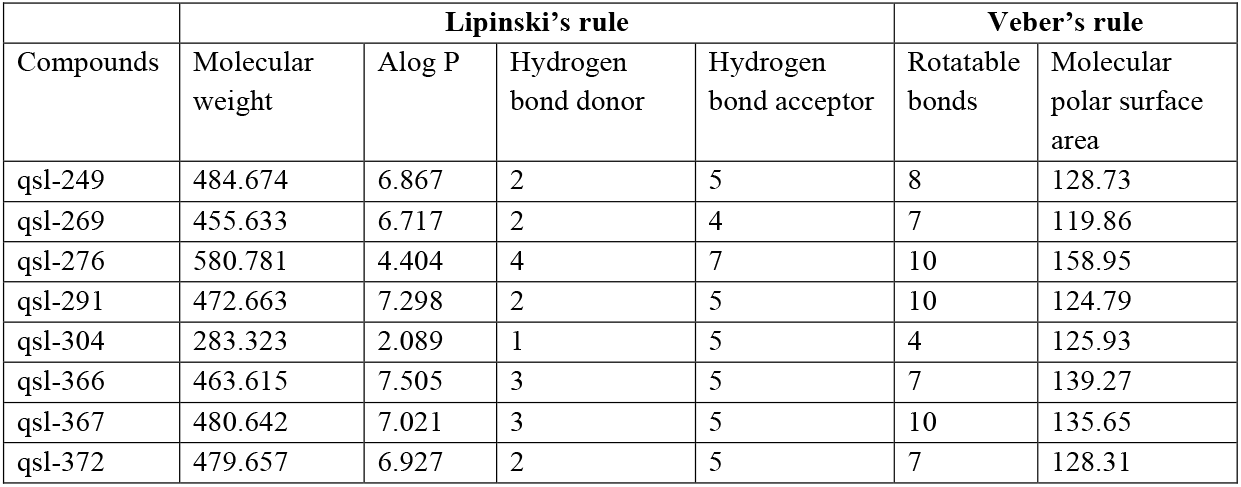
The hit molecules which passed the Lipinski’s rule of five and Veber’s rule

Table 2 summarizes the molecular docking scores for the top eight hit molecules along with key interactions. The eight hit molecules made hydrogen bonding interactions predominantly with the key active site residues namely D81, R84, G50, R144 and HOH427. Previous studies have established that the commonly found feature of ATPase inhibitors are hydrogen bond donor – acceptor motif *(27)*. This is well established for the top hit molecules. While cation – π interaction with R84 is considered as an important feature of certain ATPase inhibitors, it was not observed for the top eight hit molecules. Additionally, the stability of the interactions was evaluated by calculating the binding free energy with Prime – MMGBSA *(44).* The analysis revealed that the ***ΔG >*** −30 kcal/mol for all the eight hit molecules. Thus, further analysis was proceeded with the top hit eight molecules having the acceptable range of docking score, binding energy and interaction with key residues which obeyed Lipinski’s and Veber rule.

**Table 2:** Molecular Docking Scores of top eight hit molecules

| S.No. | Molecule | GScore | MMGBSA<br>$\Delta G$ bind<br>(kcal/mol) | Interacting<br>Residues -<br>Hydrogen<br>bond | Lipophilic<br>EvdW | HBond | Electro | Sitemap |
| --- | --- | --- | --- | --- | --- | --- | --- | --- |
| 1 | qsl-249 | -3.826 | -39.49 | ARG144 | -3.87 | -0.35 | -0.23 | -0.28 |
| 2 | qsl-269 | -3.711 | -38.71 | ASP57,<br>H2O | -2.99 | -1.32 | -0.43 | 0 |
| 3 | qsl-276 | -5.613 | -57.29 | H2O | -3.46 | -1.78 | -0.37 | -0.88 |
| 4 | qsl-291 | -4.096 | -46.21 | GLU50,<br>H2O | -3.26 | -1.03 | -0.33 | 0 |
| 5 | qsl-304 | -4.911 | -40.49 | ASP81,<br>ARG84,<br>H2O | -1.83 | -1.33 | -0.62 | -0.98 |
| 6 | qsl-366 | -4.988 | -49.49 |  | -4.81 | -0.66 | -0.04 | -0.28 |
| 7 | qsl-367 | -4.223 | -35.28 | ASP57,<br>ASN54, | -3.81 | -0.83 | -0.06 | -0.19 |
| 8 | qsl-372 | -4.073 | -37.44 | ASP81 | -3.84 | -0.35 | -0.35 | -0.39 |
| 9 | Novobiocin | -4.724 | -38.11 | ASN54,<br>GLU50 | -3.9 | -1.7 | -0.6 | 0 |

GScore - Total Glide Score; sum of XP terms. LipophilicEvdW - Lipophilic term derived from hydrophobic grid potential at the hydrophobic ligand atoms. HBond - ChemScore H-bond pair term. Electro - Electrostatic rewards; includes Coulomb and metal terms. Sitemap - Site Map ligand-receptor non-H bonding polar-hydrophobic terms. $\pi$ Cat - Reward for pi-cation interactions.

As a final screening process, the pharmacokinetics properties of the selected eight hit molecules were performed. Figure 4 shows a biplot figure of the 95% and 99% confidence ellipses of the blood-brain barrier (BBB) and human intestinal absorption (HIA) properties. Only one of the hit molecules, qsl-304 was within the acceptable ranges of both BBB and HIA. Table 3 depicts the detailed ADME descriptors considered. The hydrophilicity was evaluated by calculating A log P value. The values of less than 5 indicate better permeation. In this regard, the hit molecule qsl-304 showed A log P value of 2.089 depicting better hydrophilicity and permeation. Polar surface area (PSA) describes the oral bioavailability, which should be < 140. All the eight hit molecules were less than the allowed values ranging between 60 - 80. Human intestinal absorption was good only for qsl-304, whereas the other hits had either poor or extremely poor absorption values. All the eight hit molecules could be bound to plasma protein whereas none of them could penetrate BBB and not inhibit cytochrome p450. Hence all the hit compounds can easily undergo oxidation and hydroxylation during the first phase of metabolism. Among the eight hit molecules, four (qsl-269, qsl-276, qsl-304 and qsl-372) were non-hepatotoxic and three (qsl-269, qsl-304, qsl-372) were non-DTP toxic. All the eight hit molecules are non - carcinogenic, non - mutagenic and four hit molecules (qsl-249, qsl-269, qsl-291, qsl-304) were degradable. Considering all these properties qsl-304 is selected as an ideal candidate for the drug development process. When compared to the positive control, novobiocin (−38.11 kcal/mol), qsl-304 (−40.49 kcal/mol) has enhanced binding energy towards ATP binding site of *S. aureus* GyrB.

**Figure 4:**
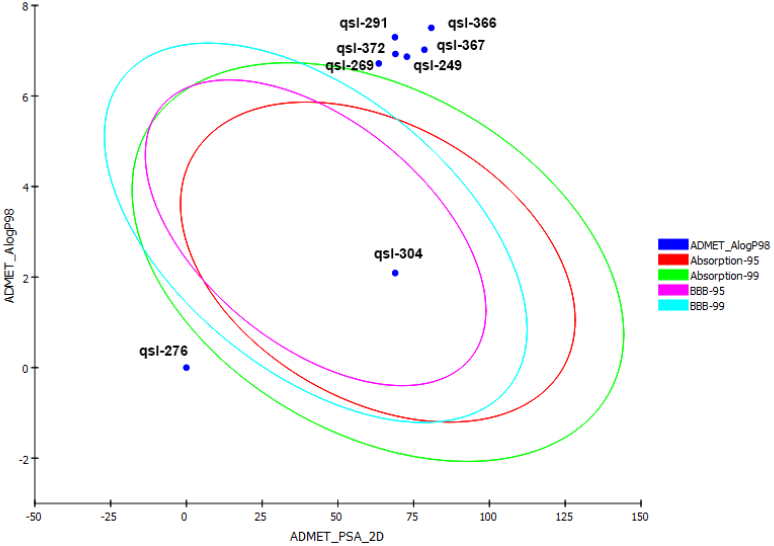
ADME/T evaluation plot of PSA vs A log P for the eight hit molecules. 99% and 95% confidence limits correspond to intestinal absorption and blood brain barrier models.

**Table 3:** In silico ADME/T prediction of the hit molecules which passed Lipinski’s rule and Veber’s rule of Drug likeliness

| ADME/T<br>PARAMETERS | qsl-249 | qsl-269 | qsl-276 | qsl-291 | qsl-304 | qsl-366 | qsl-367 | qsl-372 |
| --- | --- | --- | --- | --- | --- | --- | --- | --- |
| A log p98 | 6.867 | 6.717 | 6.524 | 7.298 | 2.089 | 7.505 | 7.021 | 6.927 |
| PSA | 72.776 | 63.48 |  | 68.829 | 68.896 | 80.808 | 78.535 | 68.896 |
| Aqueous solubility | 1 | 1 | 1 | 1 | 3 | 0 | 1 | 1 |
| HIA | 2 | 2 |  | 3 | 0 | 3 | 3 | 2 |
| PPB | 0 | 0 | 0 | 0 | 0 | 0 | 0 | 0 |
| BBB penetration | 4 | 4 |  | 4 | 3 | 4 | 4 | 4 |
| Cyp450 2D6<br>binding | 0 | 0 | 0 | 0 | 0 | 0 | 0 | 0 |
| Hepatotoxicity | Toxic | Non-Toxic | Non-Toxic | Toxic | Non-Toxic | Toxic | Toxic | Non-Toxic |
| DTP | Toxic | Non-Toxic | Toxic | Toxic | Non-Toxic | Toxic | Toxic | Non-Toxic |
| FDA rodent<br>carcinogenicity | Non-<br>carcinogenic | Non-<br>carcinogenic | Non-<br>carcinogenic | Non-<br>carcinogenic | Non-<br>carcinogenic | Non-<br>carcinogenic | Non-<br>carcinogenic | Non-<br>carcinogenic |
| Ames mutagenicity | Non-<br>mutagen | Non-<br>mutagen | Non-<br>mutagen | Non-<br>mutagen | Non-<br>mutagen | Non-<br>mutagen | Non-<br>mutagen | Non-<br>mutagen |
| Aerobic<br>biodegradability | Degradable | Degradable | Non-Degradable | Degradable | Degradable | Non-Degradable | Non-Degradable | Non-Degradable |
| Skin sensitization | None | Weak | Weak | Weak | Weak | None | None | Weak |
| Skin irritating | Mild | Mild | Mild | Mild | Mild | None | Mild | Mild |

A log P98 (atom-based log P) ( $\leq 2.0$ or $\geq 7.0$ : very low absorption). PSA (polar surface area) ( $>150$ : very low absorption). Level of aqueous solubility predicted: 0 (extremely low), 1 (very low, but possible), 2 (low), 3 (good), 4 (optimal), 5 (too soluble), 6 (warning: molecules with one or more unknown A log P calculations). HIA (human intestinal absorption), level of human intestinal absorption prediction: 0 (good), 1 (moderate), 2 (poor), 3 (very poor). PPB, plasma protein binding. f BBB (blood brain barrier), level blood brain barrier penetration prediction: 0 (very high penetrate), 1 (high), 2 (medium), 3 (low), 4 (undefined). Prediction cytochrome P4502D6 enzyme inhibition (0: non- inhibitor; 1: inhibitor). h DTP, development toxicity potential. FDA- Food, and drug administration.

Thus, this systematic screening process identified the *de novo* ligand designed qsl-304 having a higher probability of easy synthesis that makes crucial interactions for significant GyrB inhibition and finally shows desirable druggable and pharmacokinetics characteristics. It should be noted that qsl-304 makes hydrogen – bond network with Asp81 and conserved water molecule through the rhodanine moiety and the developed derivative makes hydrogen bond interaction with Arg84 unlike other protein inhibitors that make cation π stacking with Arg84 (Figure 5).

**Figure 5:**
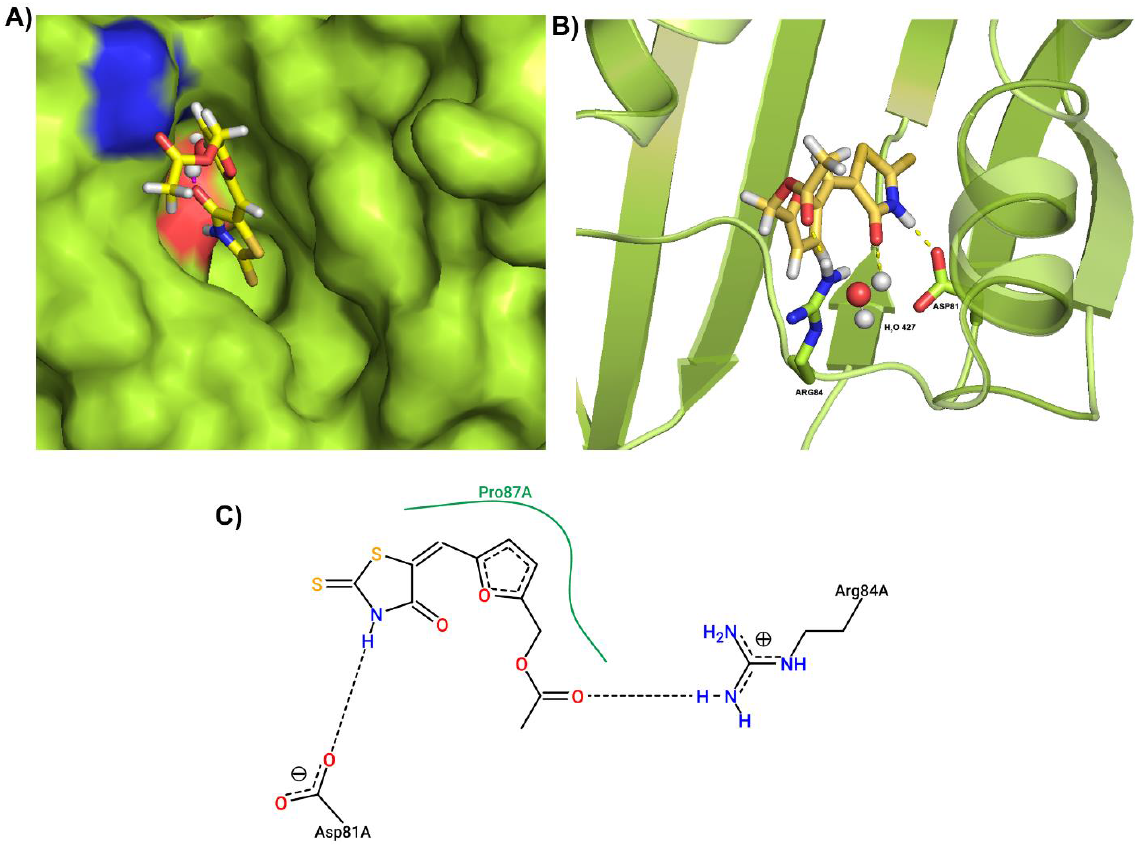
Docking pose of qsl-304 in Gyrase B ATP domain (a) GyrB is represented as green surface and docked pose of qsl-304 is represented as yellow sticks. (b) GyrB is represented as green ribbon. Active site amino acids were represented as green sticks. The binding pose of qsl-304 is represented as yellow sticks. Hydrogen bonding interaction with the active site residues and the crystallographic water molecules is shown as yellow dotted lines. C) 2D-interaction map of GyrB-qsl-304

The top-hit rhodamine scaffold hit compound (qsl-304) was synthesized by the scheme, as shown in Figure 6. The final compound was obtained as the yellow solid (1.4g, 27.5% yield): mp 146 - 147 °C. The structural details of the synthesized compound were confirmed by ^1^H NMR. The peak assignment of the spectrum are as follows: 1H NMR (300 MHz, CDCl3) δ 2.11 (s, 3H), 5.10 (s, 2H), 6.56 (d, J = 2.8 Hz, 1H), 6.78 (d, J = 2.0 Hz, 1H), 7.32 (d, J = 3.2 Hz, 1H); ^13^C NMR (75 MHz, CDCl3) δ 20.7, 57.9, 113.8, 117.6, 119.3, 124.3, 150.1, 154.1,169.4, 170.8, 195.9; LCMS obsd 284.0053, calcd 284.0056 (M + H; M = C11H10NO4S2).

**Figure 6:**
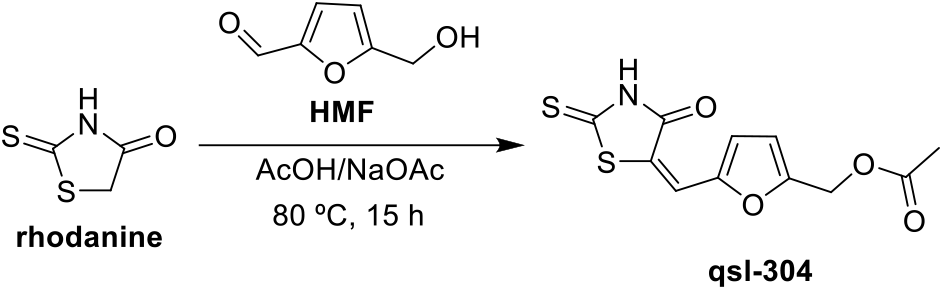
Chemical Synthesis Schema

### Biological Efficacy of qsl-304

With time, MRSA has acquired cross-resistance to most classes of antibiotics, thereby increasing the morbidity and mortality rate in hospital-acquired infections *(7)*. In this accordance, the natural existence of MRSA with cross-resistance to a failed drug, novobiocin is to be addressed. So, we initially examined the novobiocin profile of the standard strain, *S. aureus* ATCC43300 and the clinical isolates, *Sa*-P1934, *Sa*-P1920, *Sa*-P1996, *Sa*-P2003, *Sa*-2052, *Sa*-AB77, *Sa*-AB459, *Sa-* AB472 (Table S1). Among the considered clinical isolates, novobiocin resistance was observed for *sa*-P1920, *sa*-P1996, *sa*-P1920, *sa*-P1996, *sa*-P1934, *sa*-P2003, *sa*-AB459 and *sa*-AB77. The isolates *sa*-P2052, *sa*-P2040 and *sa*-AB472 were susceptible to novobiocin, whereas it showed a moderate effect on the standard strain, ATCC 43300. For the current study, the novobiocin resistant clinical strain, *sa*-P2003 was chosen.

Dose-response (500-3.9μg/mL) effect of the hit molecule, qsl-304 with rhodanine scaffold was examined in the novobiocin resistant/intermediate *S. aureus* strains (*sa*-P2003 and ATCC 43300) (Figure 7). The minimum inhibitory concentration (IC50) of qsl-304 against ATCC43300 and *sa-* P200 was observed to be 6.97 and 31.23 μg/mL respectively, indicating strain-dependent effects. Several studies have demonstrated the antibacterial efficacy of rhodanine derived molecules against both Gram-positive and Gram-negative pathogens *(20).* The presence of rhodanine scaffold efficiently enhances the antibacterial activity through peptidoglycan inhibition and prevents critical metabolic processes *(45).* Studies have also shown the targeted action on DNA gyrase of *E. coli (22)* and *S. aureus (21,39).*

**Figure 7.**
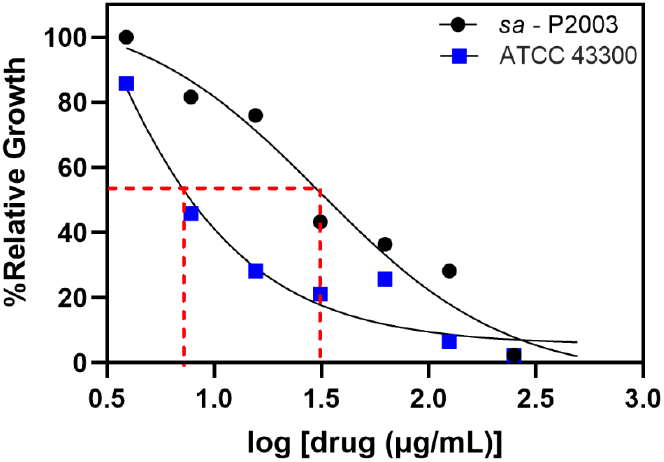
Dose Response Curve of qsl-304 against ATCC43300 (IC50 =6.97μg/mL) and sa-P2003 (IC50 =31.23μg/mL)

The bactericidal evaluation of qsl304 was tested, and the results are shown in Table 4. The bactericidal activity is defined when the initial inoculum is reduced by 3log10 CFU/mL. In the current study, the bactericidal effect was not observed until 16x MIC indicating that qsl-304 is bacteriostatic. A maximum of only ~1.5 log reduction was observed in the 16x MIC treatment. While bactericidal action is preferred in most of the cases, bacteriostatic antibiotics (clindamycin) were shown to overcome toxic shock syndrome in *S. aureus (46)*. Further studies are underway to explore the extended activities of qsl-304. Time-dependent comparative analysis on the qsl-304 treated and the untreated *S. aureus* cells showed delayed killing kinetics for the qsl-304 treatment extending generation time to 4. 25 h when compared to the control having 2.11 h (Figure 8).

**Figure 8:**
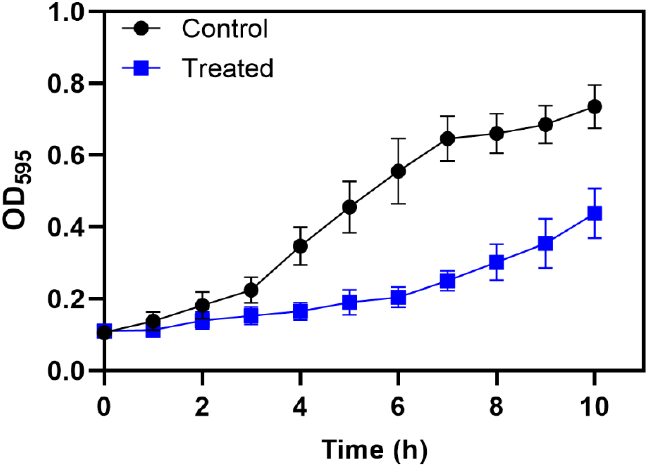
Time-Kill studies of qsl304 with Sa-P2003 having the extended generation time (4.25 h) as compared to the control (2.11 h)

**Table 4:** Time Kill Kinetics of qsl-304 antimicrobial activity at higher concentrations. <3log_10_ fold difference denotes bacteriostatic effect.

|  | Log10 CFU difference |  |  |  |  |  |
| --- | --- | --- | --- | --- | --- | --- |
| Time (h) | 0.5X MIC | 1X MIC | 2X MIC | 4X MIC | 8X MIC | 16X MIC |
| 2 | -0.13109 | -0.11522 | -0.08318 | -0.18986 | -0.3181 | -1.09293 |
| 4 | -0.18154 | -0.32239 | -0.28951 | -0.49282 | -0.65238 | -1.38441 |
| 6 | -0.07488 | -0.43306 | -0.40239 | -0.6479 | -0.78419 | -1.54983 |
| 8 | -0.00485 | -0.34024 | -0.47321 | -0.71939 | -0.89034 | -1.66343 |
| 10 | -0.00675 | -0.22468 | -0.41521 | -0.63519 | -0.7142 | -1.54426 |
| 24 | -0.08665 | -0.1568 | -0.38371 | -0.65963 | -0.8006 | -1.62342 |

The frequency of spontaneous resistance of qsl-304 was observed for the considered strains ATCC43300 and *sa*-P2003. The data was significant to show a gradual decrease in the colony forming unit (CFU/ml) on exposure to higher doses (0.5x to 4xMIC) of qsl-304 (Table 5). 4xMIC of qsl304 showed a 60% decrease in cell counts when compared to 0.5xMIC. Similar effects of qsl-304 were observed in both the strains confirming no cross-resistance. The data was comparable with the recent work of GlaxoSmithKline (GSK), wherein a series of DNA gyrase inhibitors were identified that invariably developed no cross-resistance with the current topoisomerase -targeting antibacterial agents *(47).*

**Table 5:**
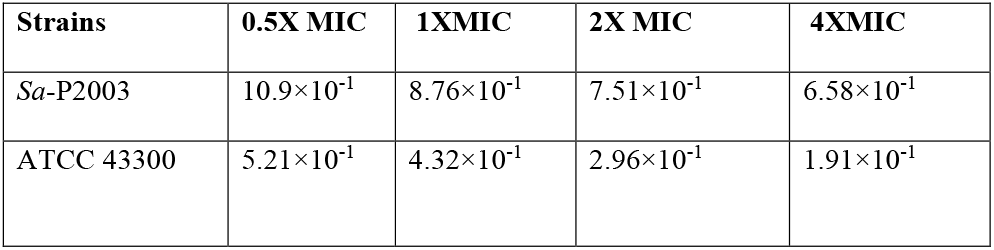
Frequency of spontaneous mutation (CFU/mL)

| Strains | 0.5X MIC | 1X MIC | 2X MIC | 4X MIC |
| --- | --- | --- | --- | --- |
| *Sa*-P2003 | $10.9 \times 10^{-1}$ | $8.76 \times 10^{-1}$ | $7.51 \times 10^{-1}$ | $6.58 \times 10^{-1}$ |
| ATCC 43300 | $5.21 \times 10^{-1}$ | $4.32 \times 10^{-1}$ | $2.96 \times 10^{-1}$ | $1.91 \times 10^{-1}$ |

While previous studies *(20)* were conducted with the standard strains of the MRSA/MSSA, the current study was extended to the clinical isolate of novobiocin resistance strain as well the cross-resistant reference strain ATCC43300. Besides, we showed that the hit compound is less prone to the development of resistance. Further studies on the effect of supercoiling and reduction in the virulence characteristics are currently in progress.

## Supporting information

Mass Spectra Data

## Acknowledgements

**AP** wish to express her gratitude to SASTRA Deemed to be University for providing her an opportunity to carry out the present work as a part of her Post-graduate dissertation work. **SV** wishes to express her sincere thanks to DST-INSPIRE (IF170369) for the financial support. **AP, SV, KS, AS, VP, PN and SAP** acknowledge SASTRA Deemed to be University, Thanjavur for extending support to use Schrödinger software suite for carrying out *in silico* studies and Professor Sumana (Department of Microbiology, JSS Medical College and JSS University, India) for the clinical *S. aureus* strains.

## Author Contributions

Study conception design and project administration: **KS**, **PN** and **SAP**; Computational data analysis and interpretation of results: **AP**, **AS** and **KS**; qsl-304 synthesis and characterization: **AP** and **VP**; *in vitro* data acquiring: **AP**; *in vitro* data analysis and interpretation of results: **AP**, **SV**, **PN** and **SAP**; Original draft manuscript preparation, writing, reviewing and editing: **AP**, **SV**, **KS**, **PN** and **SAP**. All authors have read and agreed to published version of the manuscript.

## Transparency declaration

The authors declare that the research was conducted in the absence of any commercial or financial relationships that could be construed as a potential conflict of interest.

---

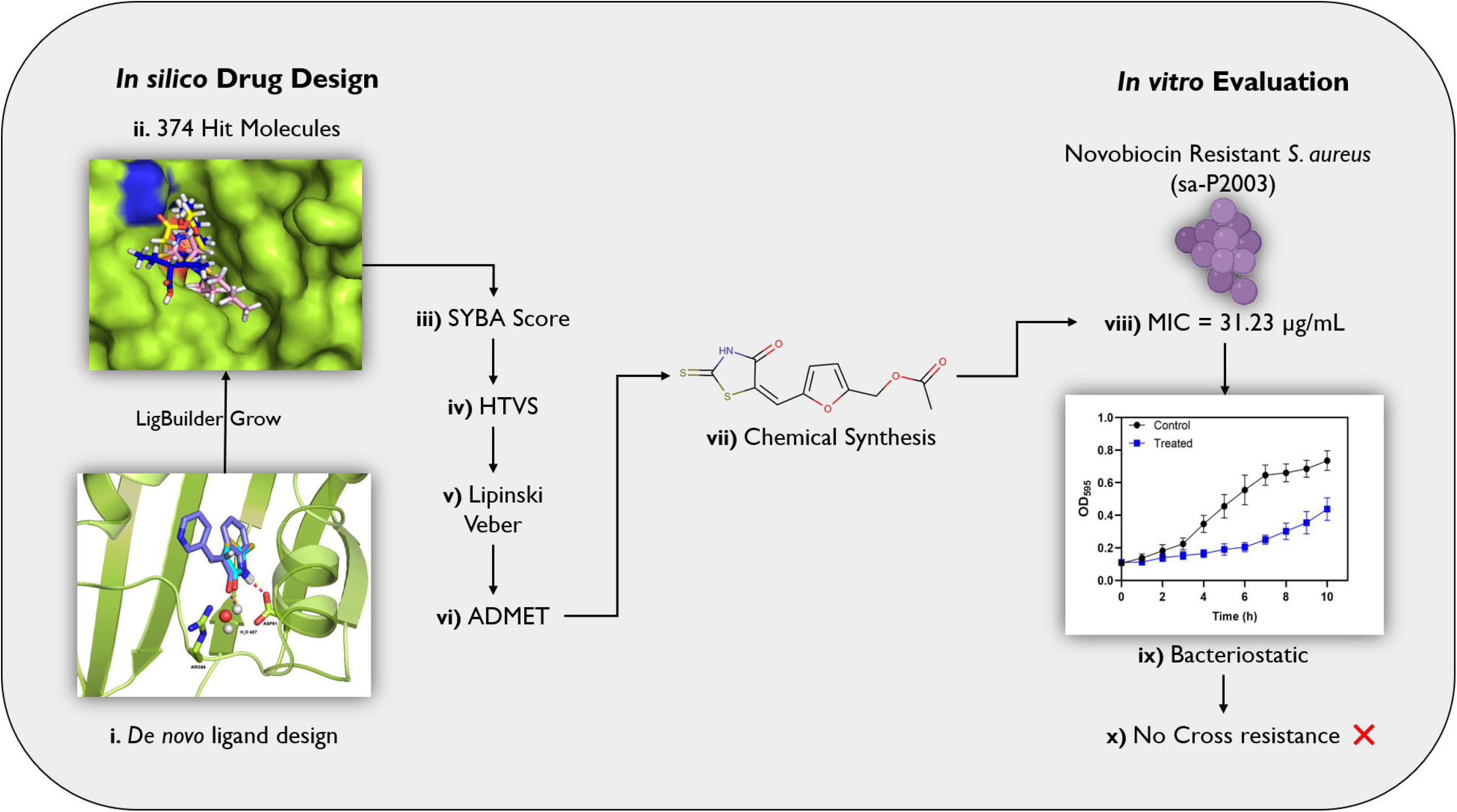

