## Supplemental Figures for "Design of a novel DNA Gyrase B inhibitor with a rhodanine scaffold: *in silico* and *in vitro* approaches"

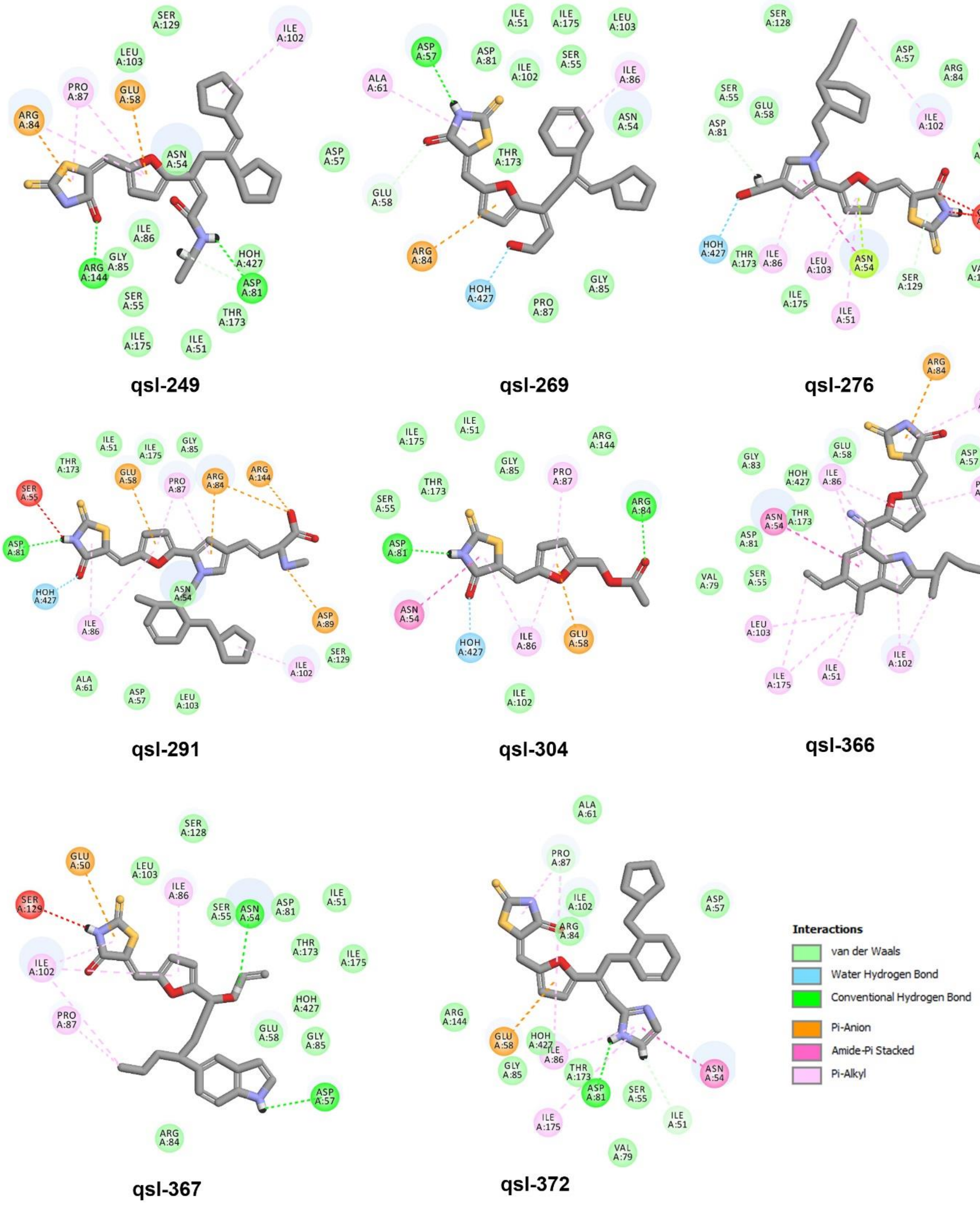


*Figure S1: 2D interaction map of the eight top hit molecules.*

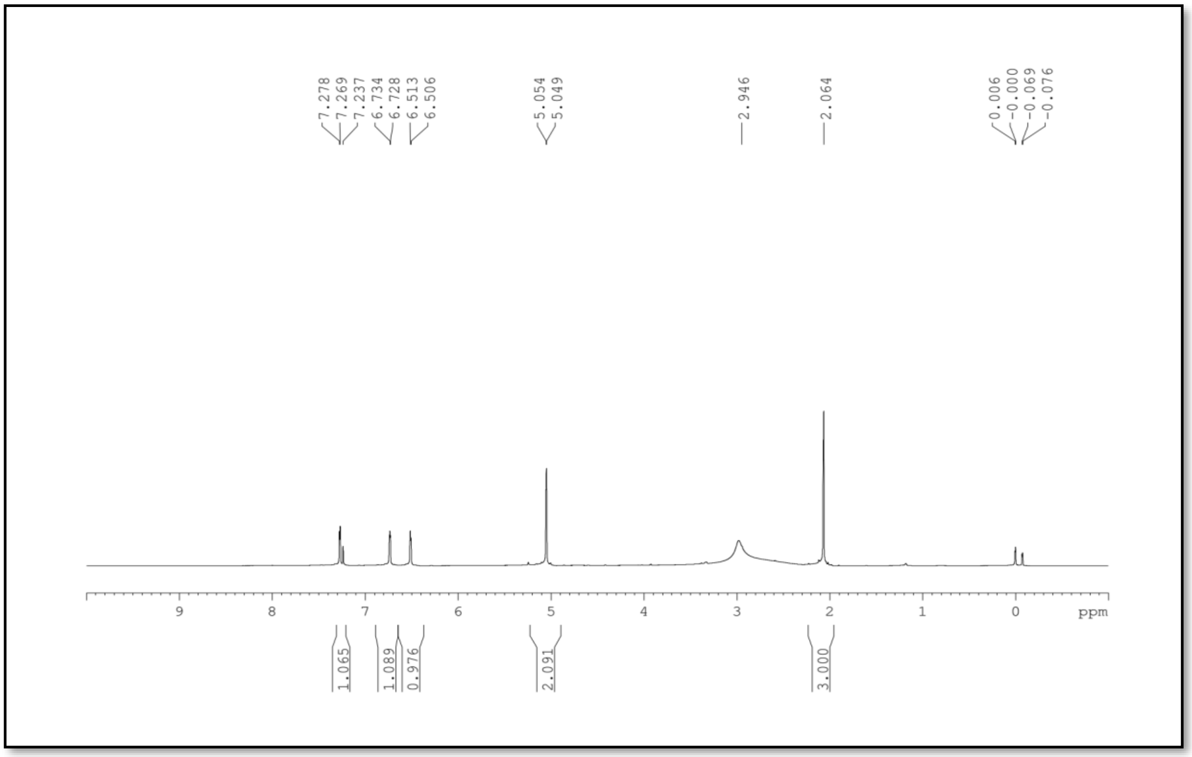


***Figure S2:*** *¹H-NMR data of qsl-304*

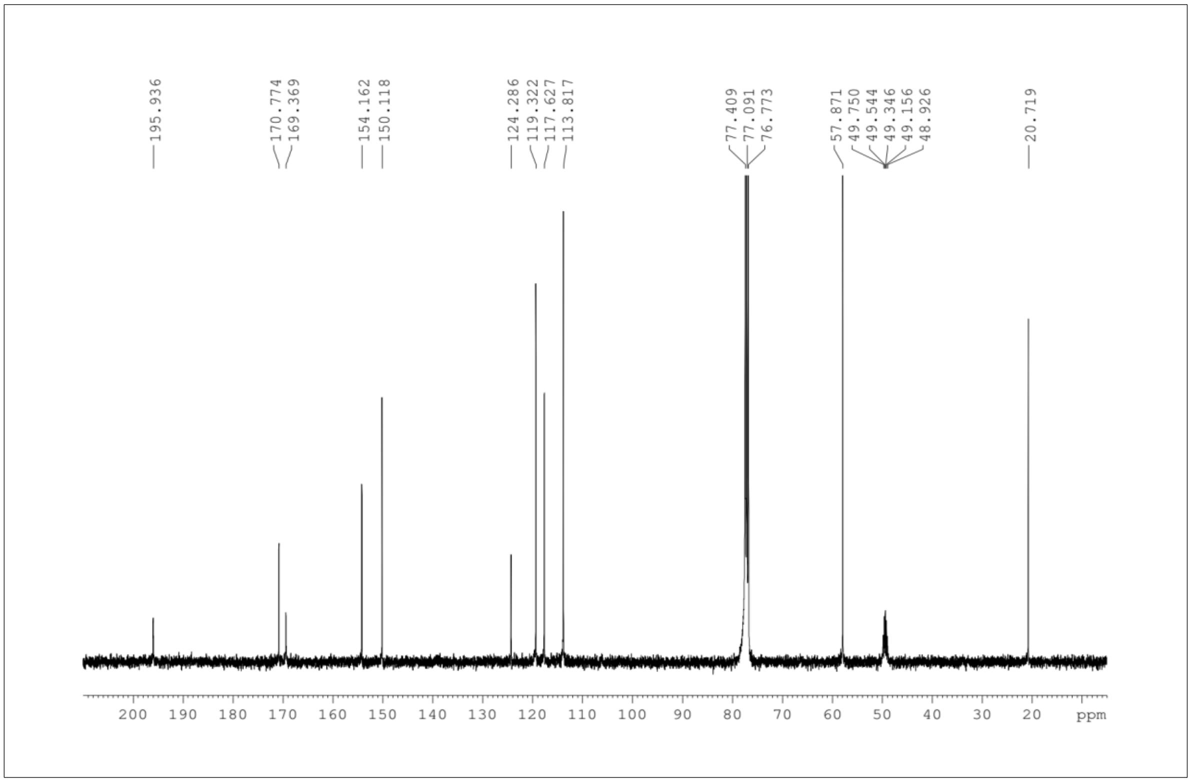


***Figure S3:*** *¹³C-NMR data of qsl-304*

***Table S1:*** *Novobiocin Resistance Profile of the clinical isolates of Staphylococcus aureus. S: Sensitive; I: Intermediate; R: Resistance*

| *S. aureus* Strain | | Profile |
|---|---|---|
| Reference | ATCC 43300 | |
| Pus Isolates | P1920 | |
| | P2052 | |
| | P1996 | |
| | P2040 | |
| | P1934 | |
| | P2003 | |
| Abscess Isolates | AB472 | |
| | AB459 | |
| | AB77 | |

| | |
|---|---|
| S | <0.25 µg/mL |
| I | 1 µg/mL |
| R | > 2 µg/mL |
