## Supplementary material for "Design of a novel DNA Gyrase B inhibitor with a rhodanine scaffold: *in silico* and *in vitro* approaches": Mass Spectra Data

### Analysis Report

Agilent Trusted Answers

#### Sample Information

| | | | |
|---|---|---|---|
| **Name** | QSL304 | **Data File Path** | D:\MassHunter\Data\OTHERS\SASTRA DEEMED UNIVERSITY\QSL304.d |
| **Sample ID** | | **Acq. Time (Local)** | 11-11-2020 15:24:37 (UTC+05:30) |
| **Instrument** | Instrument 1 | **Method Path (Acq)** | D:\MassHunter\Methods\TRAINING\BOTTLE C POSITIVE.m |
| **MS Type** | QTOF | **Version (Acq SW)** | 6200 series TOF/6500 series Q-TOF B.09.00 (B9044.1 SP1) |
| **Inj. Vol. (ul)** | 4 | **IRM Status** | Success |
| **Position** | P2-D4 | **Method Path (DA)** | D:\MassHunter\Data\OTHERS\SASTRA DEEMED UNIVERSITY\QSL304.d\Results\Qual\Version4\ORGANIC MOLECULES.m |
| **Plate Pos.** | | **Target Source Path** | |
| **Operator** | | **Result Summary** | 1 qualified (1 targets) |

#### Compound Summary

| Cpd | Name | Formula | RT | Mass | CAS | ID Source | Score | Score (Lib) | Score (DB) | Score (MFG) | Algorithm |
|---|---|---|---|---|---|---|---|---|---|---|---|
| 1 | | C11 H9 N O4 S2 | 0.296 | 282.9978 | | FBF | 99.70 | | | | FBF |

#### Compound Details

##### Cpd. 1: C11 H9 N O4 S2

| Name | Formula | RT | RI | Mass | Score | Algorithm | Lib/DB |
|---|---|---|---|---|---|---|---|
| | C11 H9 N O4 S2 | 0.296 | | 282.9978 | 99.70 | FBF | |

| Species | m/z | Score (Lib) | Num Spectra | Score (DB) | Score (MFG) | Score (RT) |
|---|---|---|---|---|---|---|
| M+ (M+H)+ (M+NH4)+ (M+Na)+ (M+K)+ | 282.9986 284.0053 301.0315 305.9870 321.9612 | | | | | |

*Compound ID Table*

| Name | Formula | Species | RT | RT Diff | Mass | CAS | ID Source | Score | Score (DB) | Score (MFG) |
|---|---|---|---|---|---|---|---|---|---|---|
| | C11 H9 N O4 S2 | M+ (M+H)+ (M+NH4)+ (M+Na)+ (M+K)+ | 0.296 | | 282.9978 | | FBF | 99.70 | | |

*Compound Chromatograms (overlaid)*

*Structure*

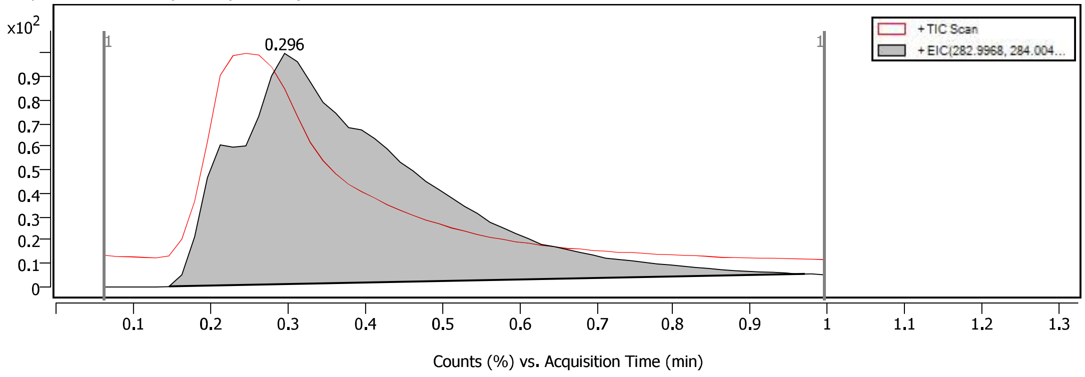


*Compound Chromatograms (overlaid)*

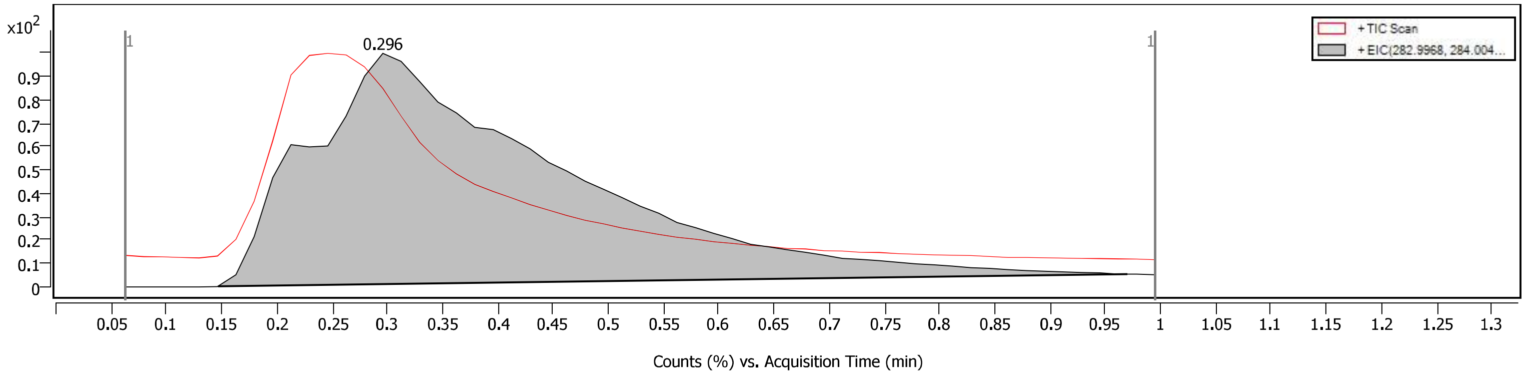

*Compound Chromatogram(s)*

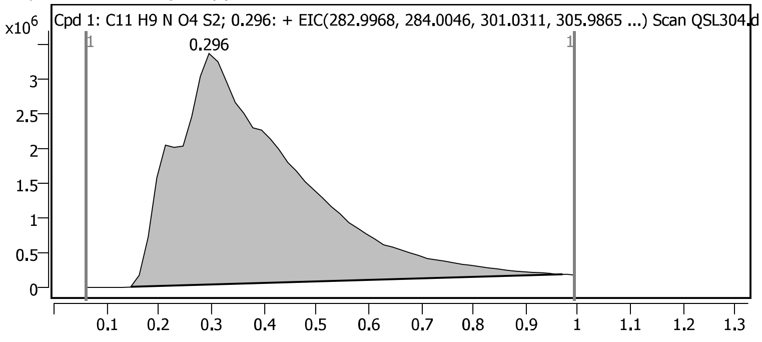


*Compound Spectra (overlaid)*

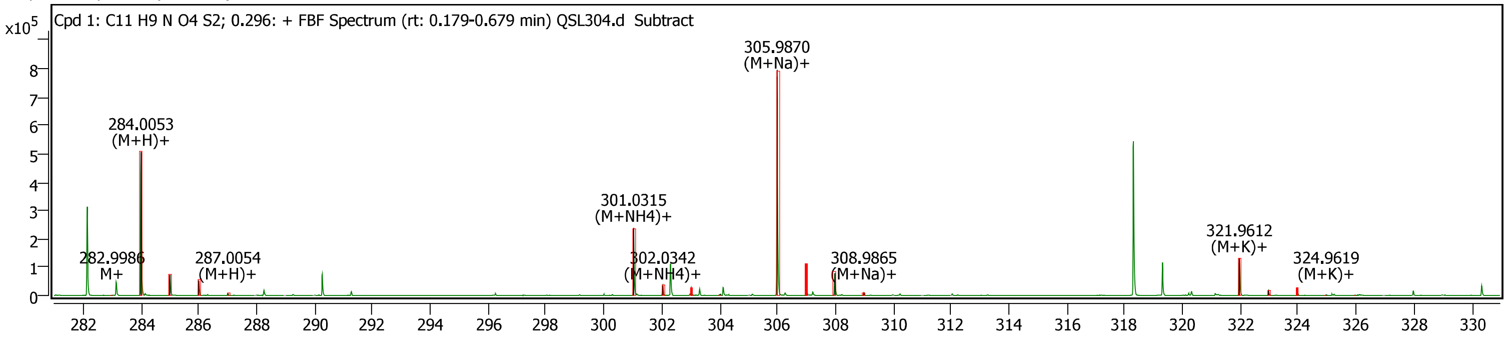


*Spectrum Peaks*

| m/z | Z | Abund | Diff (ppm) | Height % | Height % (Calc) | Ion Species | Formula |
|---|---|---|---|---|---|---|---|
| 282.9986 | 1 | 792 | 6.58 | 100.00 | 100.00 | M+ | C11H9NO4S2 |
| 284.0053 | 1 | 506721 | 2.51 | 100.00 | 100.00 | (M+H)+ | C11H9NO4S2 |
| 285.0079 | 1 | 72600 | 1.95 | 14.33 | 14.11 | (M+H)+ | C11H9NO4S2 |
| 286.0025 | 1 | 53146 | 2.33 | 10.49 | 10.69 | (M+H)+ | C11H9NO4S2 |
| 287.0054 | 1 | 7167 | 3.73 | 1.41 | 1.34 | (M+H)+ | C11H9NO4S2 |
| 301.0315 | 1 | 239188 | 1.23 | 100.00 | 100.00 | (M+NH4)+ | C11H9NO4S2 |
| 302.0342 | 1 | 34381 | 1.34 | 14.37 | 14.51 | (M+NH4)+ | C11H9NO4S2 |
| 303.0289 | 1 | 26005 | 1.43 | 10.87 | 10.75 | (M+NH4)+ | C11H9NO4S2 |
| 304.0204 | 1 | 4407 | -34.10 | 1.84 | 1.39 | (M+NH4)+ | C11H9NO4S2 |
| 305.9870 | 1 | 796220 | 1.47 | 100.00 | 100.00 | (M+Na)+ | C11H9NO4S2 |
| 306.9897 | 1 | 108300 | 1.41 | 13.60 | 14.10 | (M+Na)+ | C11H9NO4S2 |
| 307.9842 | 1 | 80026 | 1.30 | 10.05 | 10.69 | (M+Na)+ | C11H9NO4S2 |
| 308.9865 | 1 | 10429 | 0.88 | 1.31 | 1.34 | (M+Na)+ | C11H9NO4S2 |
| 321.9612 | 1 | 132050 | 2.20 | 100.00 | 100.00 | (M+K)+ | C11H9NO4S2 |
| 322.9640 | 1 | 18567 | 2.26 | 14.06 | 14.11 | (M+K)+ | C11H9NO4S2 |
| 323.9587 | 1 | 22693 | 2.01 | 17.19 | 17.91 | (M+K)+ | C11H9NO4S2 |
| 324.9619 | 1 | 3168 | 3.84 | 2.40 | 2.36 | (M+K)+ | C11H9NO4S2 |
| 325.9608 | 1 | 1752 | 14.49 | 1.33 | 1.15 | (M+K)+ | C11H9NO4S2 |

*Compound Spectra*

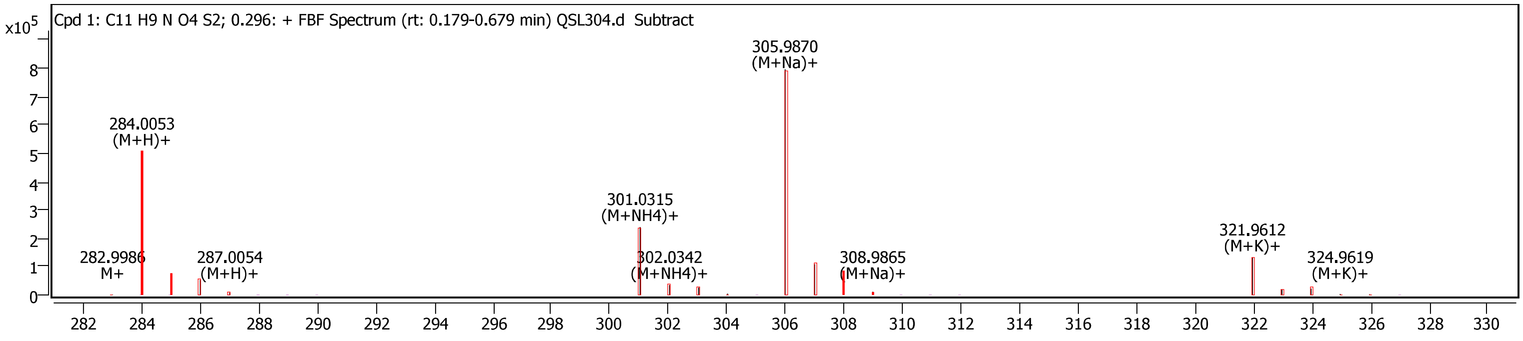
